# GnRH pulse generator frequency is modulated by kisspeptin and GABA-glutamate interactions in the posterodorsal medial amygdala in female mice

**DOI:** 10.1101/2021.11.27.470131

**Authors:** G Lass, XF Li, E Wall, RA de Burgh, D Ivanova, C McIntyre, XH Lin, WH Colledge, SL Lightman, KT O’Byrne

**Affiliations:** Department of Women and Children’s Health, Faculty of Life Sciences and Medicine, King’s College London, Guy’s Campus, SE1 1UL, UK; Reproductive Physiology Group, Department of Physiology, Development and Neuroscience, University of Cambridge, Cambridge, CB2 3EG UK; Henry Wellcome Laboratories for Integrative Neuroscience and Endocrinology, The Dorothy Hodgkin Building, University of Bristol, Whitson Street, Bristol BS1 3NY, UK

## Abstract

Kisspeptin neurons in the arcuate nucleus of the hypothalamus generate GnRH pulses, and act as critical initiators of functional gonadotrophin secretion, and reproductive competency. However, kisspeptin in other brain regions, most notably the posterodorsal subnucleus of the medial amygdala (MePD), plays a significant modulatory role over the hypothalamic kisspeptin population; our recent studies using optogenetics have shown that low frequency light stimulation of MePD kisspeptin results in increased LH pulse frequency. Nonetheless, the neurochemical pathways that underpin this regulatory function remain unknown. To study this, we have utilised an optofluid technology, precisely combining optogenetic stimulation with pharmacological receptor antagonism, to investigate the neurotransmission involved in this circuitry. We have shown that functional neurotransmission of both GABA_A_ and glutamate is a requirement for effective modulation of the GnRH pulse generator by amygdala kisspeptin neurons.

## Introduction

Recent investigations have revealed a significant stimulatory role for kisspeptin in the posterodorsal subnucleus of the medial amygdala (MePD) in GnRH pulse generator modulation. Our previous optogenetic studies showed that sustained low-frequency blue light stimulation activated these neurons to increase LH pulse frequency, a proxy for GnRH pulse generator frequency (Lass *et al*., 2020). This finding built upon our previous neuropharmacological approach using intra-MePD infusions of kisspeptin receptor (KISS1R) agonists and antagonists, which respectively increased LH secretion or decreased LH pulse frequency. (Comninos *et al*., 2016). However, the mechanisms underlying this neuronal population’s stimulatory role over pulsatile LH secretion have not been studied. Glutamate and GABA are the major stimulatory and inhibitory neurotransmitters in the mammalian brain, and many neuronal networks rely on the balance between these two to regulate their activity (Petroff, 2002). Therefore, these two neurotransmitters are sensible candidates to be used by the amygdala neuronal networks underlying the upstream, extra-hypothalamic regulation of the GnRH pulse generator. Unsurprisingly, both GABA and glutamate neurons are found in the MePD (Choi *et al*., 2005; Westberry and Meredith, 2016), and pharmacological antagonism of both has deleterious effects on several aspects of reproductive physiology (Oberlander *et al*., 2009; Polston *et al*., 2001, Li *et al*., 2015). In rats, blocking AMPA and NMDA glutamate receptors with CNQX and AP5, respectively, impedes activation of MePD neurons in response to vaginal-cervical stimulation, thereby preventing the pregnancy or pseudopregnancy response following intromission; AMPA antagonism also disrupts estrous cycles (Oberlander *et al*., 2009; Polston *et al*., 2001). Furthermore, MePD NMDA and GABA_A_R antagonism causes a weight-independent advancement of puberty (Li *et al*., 2015). Thus, GABA and glutamate play an important role in the MePD in reproductive physiology.

The interesting dichotomy between the well-known suppressive role of the MePD in reproductive physiology, and the emerging activatory function of kisspeptin within this amygdaloid subnucleus, has led to a hypothesis that MePD kisspeptinergic activity may stimulate GABAergic interneurons within the MePD that in turn synapse with, and inhibit, GABAergic projection efferents from the MePD, resulting in an overall disinhibition of the latter. Evidence to support this hypothesis stems from the knowledge that there is a significant population of GABAergic neurons that project from the MePD to reproductive neural centres such as those in the hypothalamus (Choi *et al*., 2005; Bian *et al*., 2008), and inhibitory GABA interneurons, specifically, have also been detected in this subregion (Bian, 2013; Keshavarzi *et al*., 2014). This is in line with the fact that the MePD is a pallidal subnucleus, due to its embryological origins in the caudoventral medial ganglionic eminence, indicating its similarity to other neural complexes which contain a classical GABA-GABA disinhibitory system (Pardo-Bellver *et al*., 2012). Furthermore, other subnuclei of the amygdala, such as the basolateral amygdala and posteroventral medial amygdala, have been shown to exhibit functional glutamatergic signalling onto GABA interneurons (Polepalli *et al*., 2010; Keshavarzi *et al*., 2014; Sharp, 2017), supporting the hypothesis of an alternative glutamate-GABA-GABA pathway by which kisspeptin may activate the disinhibitory system.

It is therefore critical to investigate the GABAergic and glutamatergic signalling within the MePD with respect to kisspeptin and its action over GnRH pulse generator activity. To achieve this, a dual approach of simultaneously combining optogenetic activation and pharmacological antagonists was used via the implantation of an intra-MePD optofluid cannula. By optically stimulating the kisspeptin neurons in the presence or absence of glutamate or GABA antagonists during frequent blood sampling for measurement of LH pulses, it could be determined if either of these neurotransmitters are involved in the GnRH pulse generator modulating role of MePD kisspeptin.

## Materials and methods Animals

Breeding pairs of Kiss-Cre^+/-^:tdTomato^+/+^ transgenic mice were obtained from the Department of Physiology Development and Neuroscience, University of Cambridge, UK; the Kiss-CRE mice carry a tdTomato transgene activated by CRE to label MePD Kiss1 neurons. Litters from breeding pairs were genotyped using a multiplex polymerase chain reaction (PCR) protocol to detect heterozygosity for the Kiss-Cre or wild-type allele as previously described (Yeo *et al*., 2016, Lass *et al*., 2020). Adult female mice (19-23 g), heterozygous for the *Kiss-Cre* transgene were individually housed under controlled conditions (12:12 h light-dark cycle, lights on at 07:00 h, 25±1°C) with ad libitum access to food and water. All procedures were approved by the Animal Welfare and Ethical Review Body (AWERB) Committee at King’s College London, in accordance with the United Kingdom Home Office Animals (Scientific Procedures) Act 1986.

### Stereotaxic injection of channelrhodopsin viral construct and implantation optofluid cannula

All surgical procedures were carried out under a combination of ketamine anaesthesia (Ketamidor, 100 mg/kg, i.p.; Chanelle Vet, Berkshire, UK) and xylazine (Rompun, 10 mg/kg, i.p.; Bayer, Leverkusen, Germany) under aseptic conditions. Mice were bilaterally ovariectomised (OVX) to mitigate negative feedback of endogenous estrogen on LH secretion. Stereotaxic injection of the viral construct and implantation of the brain cannula was carried out concurrently with ovariectomy. Mice were placed in a Kopf motorised stereotaxic frame (Kopf, California, USA) and procedures were carried out using a robot stereotaxic system (Neurostar, Tubingen, Germany). Following an incision of the scalp, a small hole was drilled into the skull at a location above the MePD. The stereotaxic injection coordinates used to target the MePD were obtained from the mouse brain atlas of Paxinos and Franklin (Paxinos and Franklin, 2004) (2.1 mm lateral, 1.70 mm posterior to bregma and at a depth of 5.1 mm). Using a 2-μL Hamilton micro-syringe (Esslab, Essex, UK) attached to the stereotaxic frame micro-manipulator, 0.4 μl of the ChR2 virus, AAV9-EF1a-double floxed-hChR2(H134R)-EYFP-WPRE-HGHpA (≥ 1×10^13^ vg/mL; Addgene, Massachusetts, USA) was injected unilaterally into the right MePD over 10 mins. The needle was left in position for a further 5 mins and then slowly withdrawn over 1 min. Following the injection of AAV, mice heterozygous for *Kiss-Cre* were implanted with a dual optofluid cannula (Doric Lenses, Quebec, Canada) at the same AP and ML coordinates as the virus injection site, but at a DV such that the internal cannula targets the MePD and the fibre optic cannula is situated 0.25 mm above. Dental cement (Superbond C&B kit Prestige Dental Products, Bradford UK) was used to fix the cannula in place and the skin incision was closed with a suture. Antibiotics were administered prophylactically post-surgery (Betamox, 0.15 mg/g, S.C; Norbrook Laboratories, Newry, Northern Ireland). Mice were left for 4 weeks preceding the experimental period to allow for effective opsin expression in target regions.

### Blood sampling procedure for LH measurement

Following a 1-week recovery period from surgery, the mice were handled daily to acclimatise them to the tail-tip blood sampling procedure for measurement of LH (Lass *et al*., 2020). The blood samples were processed by ELISA as reported previously (Lass *et al*., 2020). Mouse LH standard and antibody were purchased from Harbour-UCLA (California, USA) and secondary antibody (NA934) was from VWR International (Leicestershire, UK). The intra-assay and inter-assay variations were 4.6% and 10.2%, respectively.

### In vivo optogenetic stimulation of MePD kisspeptin neurons and intra-MePD infusion of bicuculline, CGP-35348 or CNQX + AP5

On the day of the experiment, the zirconia ferrule of the implanted cannula was attached via a ceramic mating sleeve to a multimode fibre optic rotary joint patch cable (Thorlabs LTD, Ely, UK) at a length that allowed for free movement of the mice in their home cage. Blue light (473 nm wavelength; 5mW) was delivered using a Grass SD9B stimulator-controlled laser (Laserglow Technologies, Toronto, Canada). Following 1 h acclimatisation, blood samples (4 µl) were collected every 5 mins for 2.5 h. The first hour of blood collection consisted of no stimulation and in the subsequent 1.5 h, Kiss-Cre mice received optic stimulation at 5 Hz (Lass *et al*., 2020).

For the neuropharmacological manipulation of GABA or glutamate receptor signalling with or without simultaneous optogenetic stimulation, mice were connected to the laser as described above, but additionally an injection cannula connected to extension tubing preloaded with drug solution was inserted into the guide cannula of the optofluid implant immediately after connection of the fibre optic cannula. Ten mins before optic stimulation, bolus administration of selective GABA_A_R (bicuculline; Sigma-Aldrich), GABA_B_R (CGP-35348; Sigma-Aldrich) or a cocktail of both NMDAR and AMPAR glutamate receptor (AP5; Tocris; CNQX; Tocris) antagonist dissolved in artificial cerebrospinal fluid (aCSF) commenced and continued over 5 mins. The concentrations for the bicuculline, CGP-35348, AP5, and CNQX boli were 20 pmol, 4.5 nmol, 1.2 nmol, and 0.5 nmol respectively. After 10 mins, immediately prior to the onset of optogenetic stimulation, continuous drug infusion commenced and continued for the remainder of the experiment (1.5 h). The total concentrations for the continuous infusion of bicuculline, CGP-35348, AP5, and CNQX were 68 pmol, 15 nmol, 2 nmol, and 1 nmol respectively. The same regimen was used in the absence of optic stimulation. As a control, Kiss-Cre mice received aCSF (0.3 μl bolus, 1 μl continuous) in the presence of 5 Hz optic stimulation following the same timeframe as the test experiments.

### Validation of AAV injection site

Upon completion of experimental procedures, the mice were sacrificed with a lethal dose of ketamine and transcardially perfused with heparinised saline for 5 mins, followed by 10 mins of ice-cold 4 % paraformaldehyde in PBS (pH 7.4) for 15 mins using a pump (Minipuls, Gilson, Villiers Le Bel, France). Their brains were rapidly collected and post-fixed sequentially at 4 °C in 15% sucrose in 4% PFA and in 30% sucrose in 1 x PBS until they sank. Brains were then snap frozen on dry ice and stored at -80 °C until processing. Coronal brain slices (30 μm thick) were sectioned using a cryostat (Bright Instrument, Luton, UK). Every third section was collected between -1.34 mm to -2.79 mm from the bregma. Sections were mounted on glass microscope slides, air-dried and cover-slipped with Prolong Antifade mounting medium (Molecular Probes, Inc, OR, USA). Only animals expressing EYFP fluorescent protein in the MePD were analysed by using Axioskop 2 Plus microscope equipped with Axiovision version 4.7 (Zeiss).

### Statistical analysis

The Dynpeak algorithm was used to establish LH pulses (Vidal *et al*., 2012). The effect of optogenetic stimulation and neuropharmacology studies was established by comparing the mean LH inter-pulse interval (IPI), from the 1 h pre stimulation or drug administration control period to the 1.5 h experimental period. On occasions where there were no LH pulses observed in the post treatment interval the IPI was given a value of 90 mins. LH pulse parameters were analysed by a two-way repeated measures ANOVA and subsequent Tukey post-hoc test. All statistics were performed using SigmaPlot (version 14). The threshold level for statistical significance was set at P < 0.05 with data presented as mean ± SEM.

## Results

### Validation of AAV injection site and cannula position

The AAV-ChR2 virus used to infect the cells in Kiss-Cre mice was tagged with fluorescent EYFP in order to be visualised under a microscope. The mean number of tdTomato-expressing kisspeptin cells in unilaterally-injected brain sections was 24.50 ± 5.20 (mean+SEM) per animal and 20.33 ± 4.50 (∼83%) of tdTomato-expressing neurons were EYFP fluorescence positive. Analysis of images acquired from coronal sectioning of the mouse brains showed that 7 out of 9 animals had successful stereotaxic injection of AAV-ChR2 virus into the MePD, and all 7 also had successful cannula implantation into the MePD. A representative example of a coronal brain section is shown in Figure 1. Dual fluorescent labelling revealed EYFP infection in tdTomato (*Kiss-Cre*-expressing neurons) cell bodies (Fig. 1G).

**FIGURE 1.**
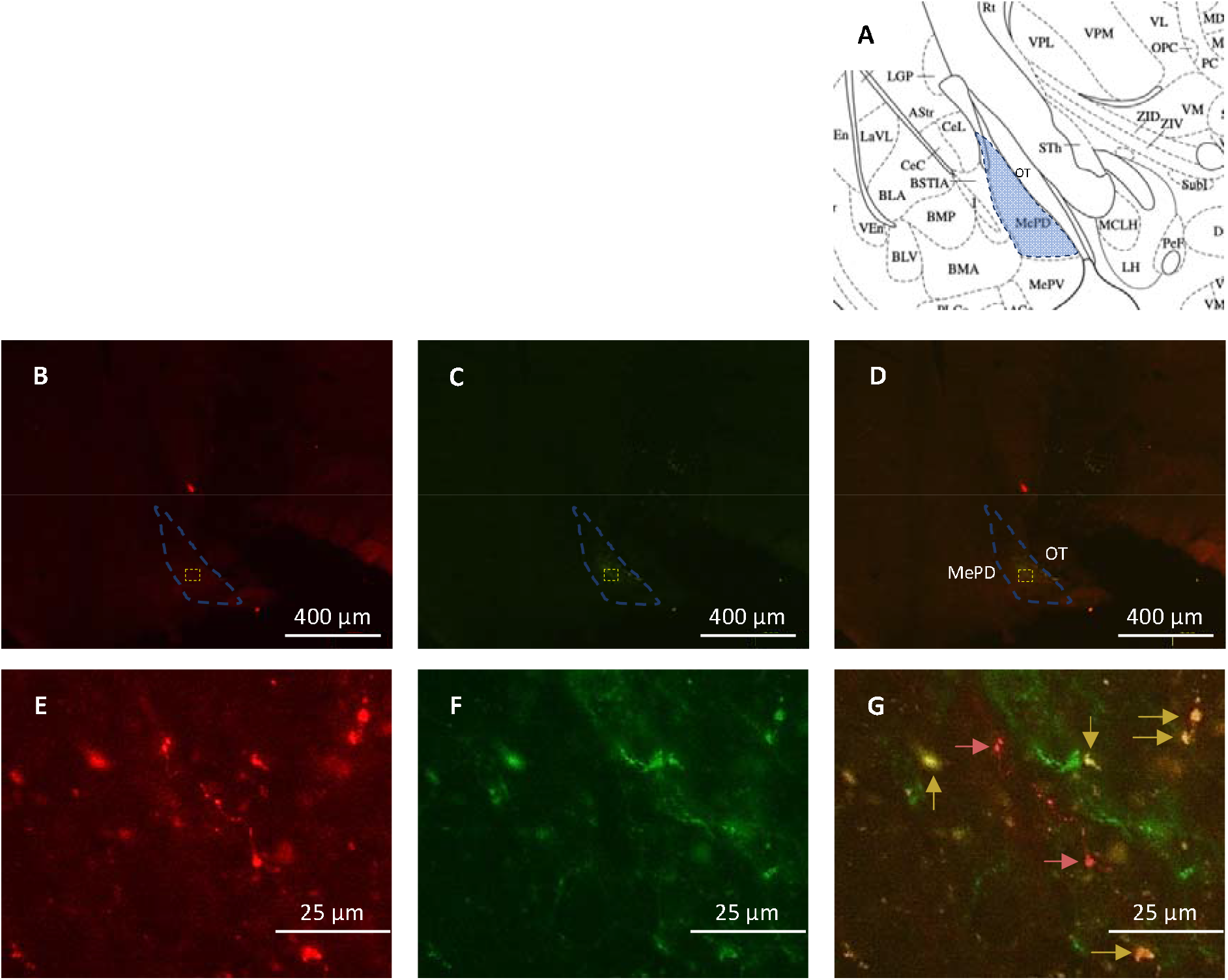
Expression of ChR2-EYFP in MePD kisspeptin neurones in Kiss-Cre mice. (A), shows a schematic representation of the MePD and its spatial relationship with the optic tract (OT). (B), shows tdTomato-expressing kisspeptin neurones, while (C) shows those cells infected with EYFP. Coronal section showing green EYFP fluorescence positive neurones in the MePD (D). (E-G), show a higher-power magnification of the area in (B-D) encased with the yellow dotted line, showing fluorescence of tdTomato (red cells), EYFP (green cells), and both (yellow cells), respectively. MePD kisspeptin neurones tagged with EYFP (labelled with yellow arrows) and not tagged with EYFP (red arrows) are shown in (G).

### Effects of sustained optical stimulation at 5 Hz, with and without a control administration of aCSF, on LH pulse frequency

After a 1 h control blood sampling period in the absence of optical stimulation, Kiss-Cre mice were stimulated at 5 Hz for 90 mins with and without administration of aCSF. In both experimental protocols, the stimulation resulted in a significant increase in LH pulse frequency (Fig. 2A and B). The mean inter-pulse interval (IPI) decreased from 25.00 ± 2.99 mins to 18.60 ± 2.07 mins (n = 7; p = 0.002) after 5 Hz stimulation only, and from 31.25 ± 1.25 mins to 22.73 ± 1.94 mins (n = 4; p = 0.002) after 5 Hz stimulation and infusion of aCSF (Fig.2C).

**FIGURE 2.**
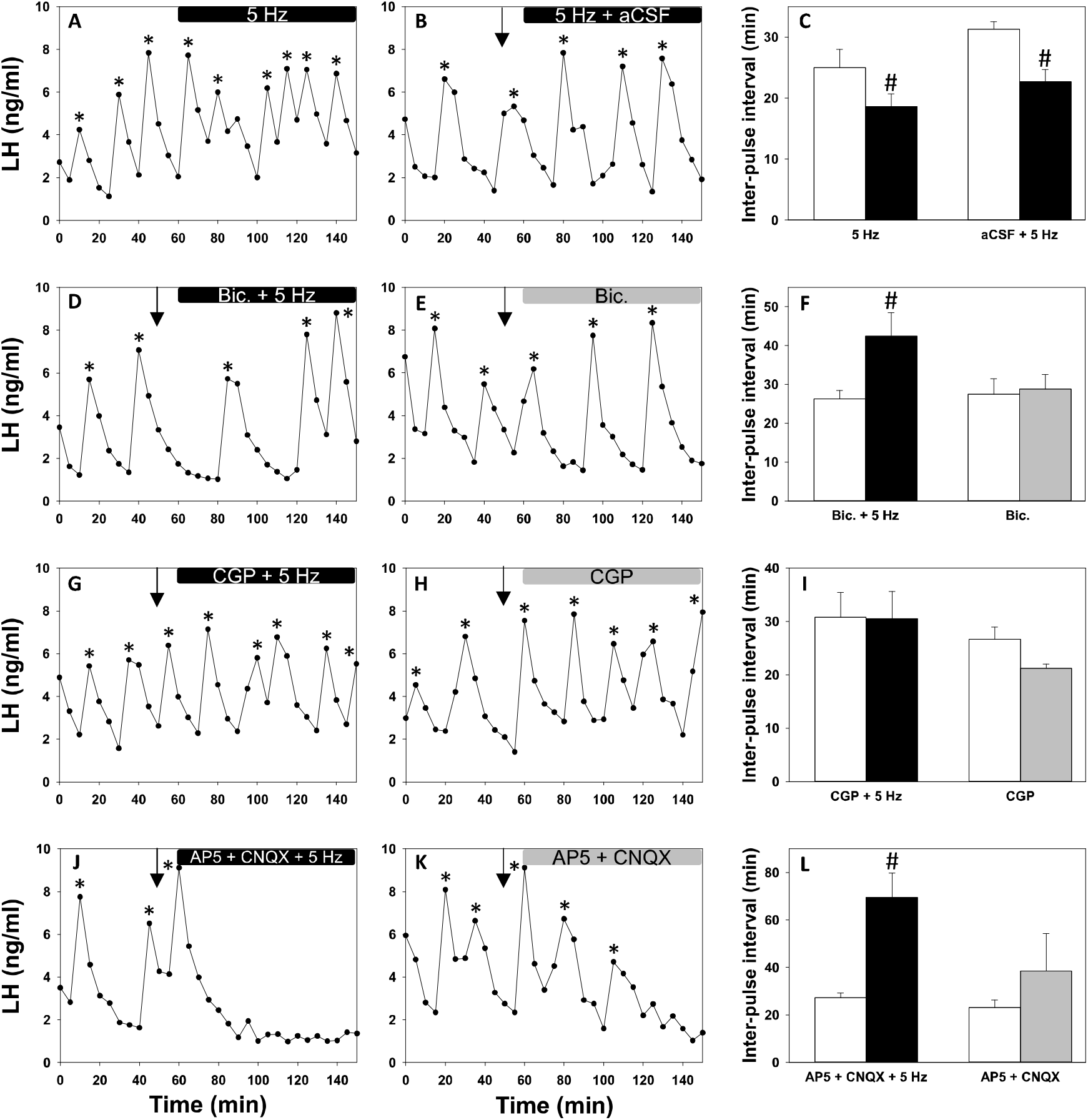
Effect of GABA and glutamate receptor antagonism with and without sustained optogenetic stimulation of MePD Kiss1 neurones on LH pulse frequency. Representative examples showing the effects of 5 Hz stimulation (A), 5 Hz + aCSF (B), bicuculline + 5 Hz (D), CGP-35348 + 5 Hz (G), AP5 + CNQX + 5 Hz (J) on pulsatile LH secretion in ovariectomised mice. Representative examples for the effects of drugs alone – bicuculline (E), CGP (H), AP5 + CNQX (K) – are also shown. Histograms showing the mean LH interpulse interval for all interventions are also provided (C, F, I, L). Sustained stimulation at 5 Hz in the presence (n = 4) and absence (n = 7) of aCSF infusion resulted in a significant reduction in the LH interpulse interval (p < 0.05). Sustained stimulation together with bicuculline (n = 5) and AP5 + CNQX (n = 6) resulted in a significant increase in LH interpulse interval. Wild type control animals did not respond to stimulation. LH pulses detected by the DynPeak algorithm are indicated with an asterisk. Bolus injections are indicated by downward black arrows. ^#^P < 0.05 vs control. Results represent the mean ± SEM.

### Effects of bicuculline, a GABA_A_R antagonist, on LH pulse frequency in the presence and absence of sustained 5 Hz optical stimulation

In the second part of the experiment, Kiss-Cre mice received a unilateral intra-MePD infusion of bicuculline with and without sustained 5 Hz optical stimulation. After the dual treatment of light and bicuculline, the average LH IPI significantly increased from 27.00 ± 2.55 mins to 42.42 ±. to 42.42 ± 7.39 mins (n = 5; p = 0.008), indicating reduced LH pulse frequency (Fig. 2D and F). There was no significant change in the IPIs before and after bicuculline administration alone, with a pre-infusion IPI of 29.00 ± 3.78 mins and a post-infusion IPI of 29.75 ± 3.78 mins (n = 5; p = 0.771) (Fig. 2E and F).

### Effects of CGP, a GABA_B_R antagonist, on LH pulse frequency in the presence and absence of sustained 5 Hz optic stimulation

In the third part of the experiment, Kiss-Cre mice received a unilateral infusion of CGP-35348, an antagonist selective for the GABA_B_R, both with and without continuous optogenetic stimulation. There was no significant difference in LH IPI between the initial 60 mins control period and subsequent period of intervention in either of these experimental protocols, however a trend towards increased LH pulse frequency during administration of CGP-35348 alone in the absence of light was observed (Fig. 2G-I). The pre- and post-intervention average IPIs for CGP-35348 with 5 Hz stimulation were 30.80 ± 4.67 mins and 30.53 ± 5.10 mins, respectively (n = 5; p = 0.954). For CGP-35348 administration alone, the pre- and post-intervention average IPIs were 26.67 ± 2.28 mins and 21.25 ± 0.72 mins, respectively (n = 5; p = 0.115).

### Effects of glutamate receptor antagonism on LH pulse frequency with and without continuous 5 Hz optic stimulation

The final protocol of the experiment involved unilateral infusion of a drug cocktail consisting of both AP5 and CNQX, antagonists for AMPA and NMDA receptors, respectively, in the presence and absence of light in Kiss-Cre mice (Fig. 2J-L). After the 60 mins control period, sustained 5 Hz optogenetic stimulation together with infusion of the antagonist resulted in a significantly decreased LH pulse frequency; indeed, in a number of cases LH pulsatility ceased altogether and the average IPI increased from 27.33 ± 1.89 mins to 69.50 ± 10.26 mins (n = 6; p = 0.007). For AP5 and CNQX alone, there was a trend of increased IPI before and after treatment from 23.04 ± 3.24 mins to 38.59 ± 15.73 mins, however this was not significant (n = 4; p = 0.253).

## Discussion

The present study highlights for the first time a potential mechanism by which kisspeptin activity in the MePD is stimulatory over the hypothalamic GnRH pulse generator. Building upon previous findings that low-frequency (5 Hz) optogenetic stimulation of MePD *Kiss1* neurons increases the frequency of pulsatile LH secretion (Lass *et al*., 2020), the results presented in this study support the hypothesis that this is reliant upon the activity of both GABA and glutamate in the MePD.

It has been shown that the MePD, with its overwhelmingly GABAergic neuronal outputs, is a significant inhibitor of gonadotrophic hormone secretion and wider facets of reproductive physiology; its stimulation and lesioning delays and advances pubertal onset, respectively, and the MePD’s activation during stress is deleterious on reproductive function and behaviour (Elwers and Critchlow, 1960; Bar-Sela and Critchlow, 1966; Lin *et al*, 2011). However, the recent findings that MePD *Kiss1* neurons stimulate the GnRH pulse generator raises the hypothesis for intranuclear GABA-GABA disinhibitory interactions, typical of limbic pallidal structures such as the MePD. The current study tested this utilising specific optogenetic activation of MePD kisspeptin neurons in conjunction with bicuculline and CGP-35348 – intra-MePD antagonists for GABA_A_R and GABA_B_R, respectively. Indeed, either of these drugs together with optic stimulation prevented the increase in LH pulse frequency seen with 5 Hz stimulation alone. The result of intra-MePD infusion of bicuculline together with 5 Hz optogenetic stimulation is a surprising one and poses an interesting question: how can intra-MePD GABA_A_R antagonism not only prevent the stimulatory effects of optogenetic stimulation, but in fact cause the opposite result of significantly reducing the activity of the GnRH pulse generator, while bicuculline alone had no effect on LH pulse frequency? We suggest a neuronal circuit, involving glutamatergic synaptic mechanisms, that may underly this phenomenon (Figure 3). Indeed, a stimulatory effect of the MePD kisspeptinergic system over the GnRH pulse generator has been linked to glutamatergic activation; the pubertal transition is tightly correlated with a developmental increase in the expression of both kisspeptin (Stephens *et al*., 2016) and glutamate (Cooke, 2011) in the MePD, and 29% of MePD *Kiss1* neurons in adult male mice coexpress vesicular glutamate transporter 2 (VGlut2) mRNA (Aggarwal *et al*., 2019); indeed, the use of glutamate as a neurotransmitter by *Kiss1* neurons has been found for the KNDy population (Voliotis *et al*., 2021) including their regulation of *Kiss1* neurons in the AVPV (Qiu *et al*., 2018). The present finding of an almost-complete ablation of pulsatile LH secretion following MePD kisspeptin activation combined with infusion of intranuclear glutamate antagonists provides further support to MePD kisspeptin effect’s dependence on glutamate. Therefore, the following mechanism is proposed: optical stimulation of MePD kisspeptin in the presence of GABA_A_R antagonists decreases LH pulse frequency by glutamatergically activating the hypothetical GABAergic projections from the MePD to KNDy neurons; whether this occurs via glutamate secretion from MePD kisspeptin cells themselves is unknown. In other words, cancelling the ability of GABA interneurons to take part in the disinhibition during optical stimulation shifts the balance from a stimulatory to inhibitory output from the MePD.

**FIGURE 3.**
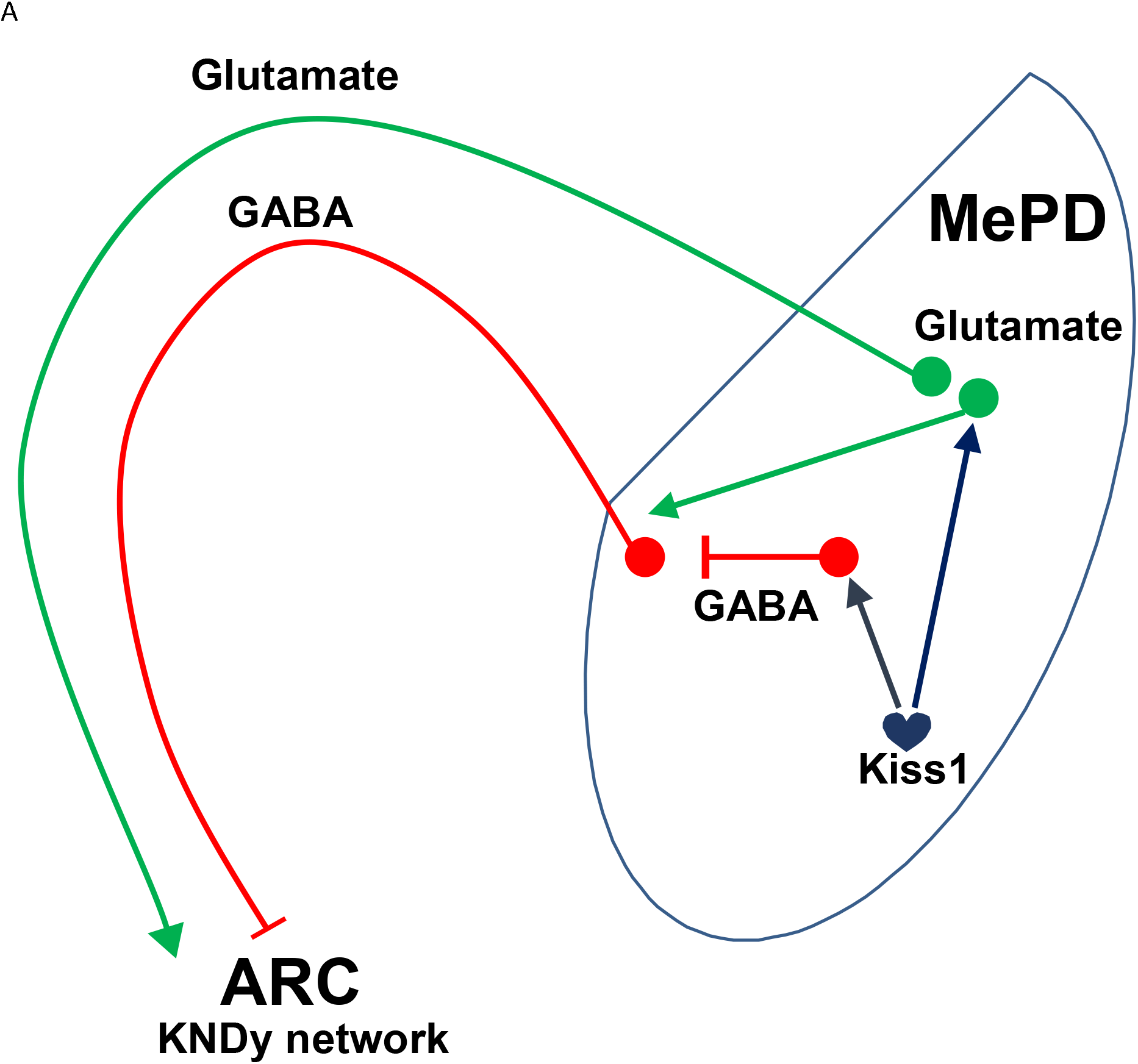
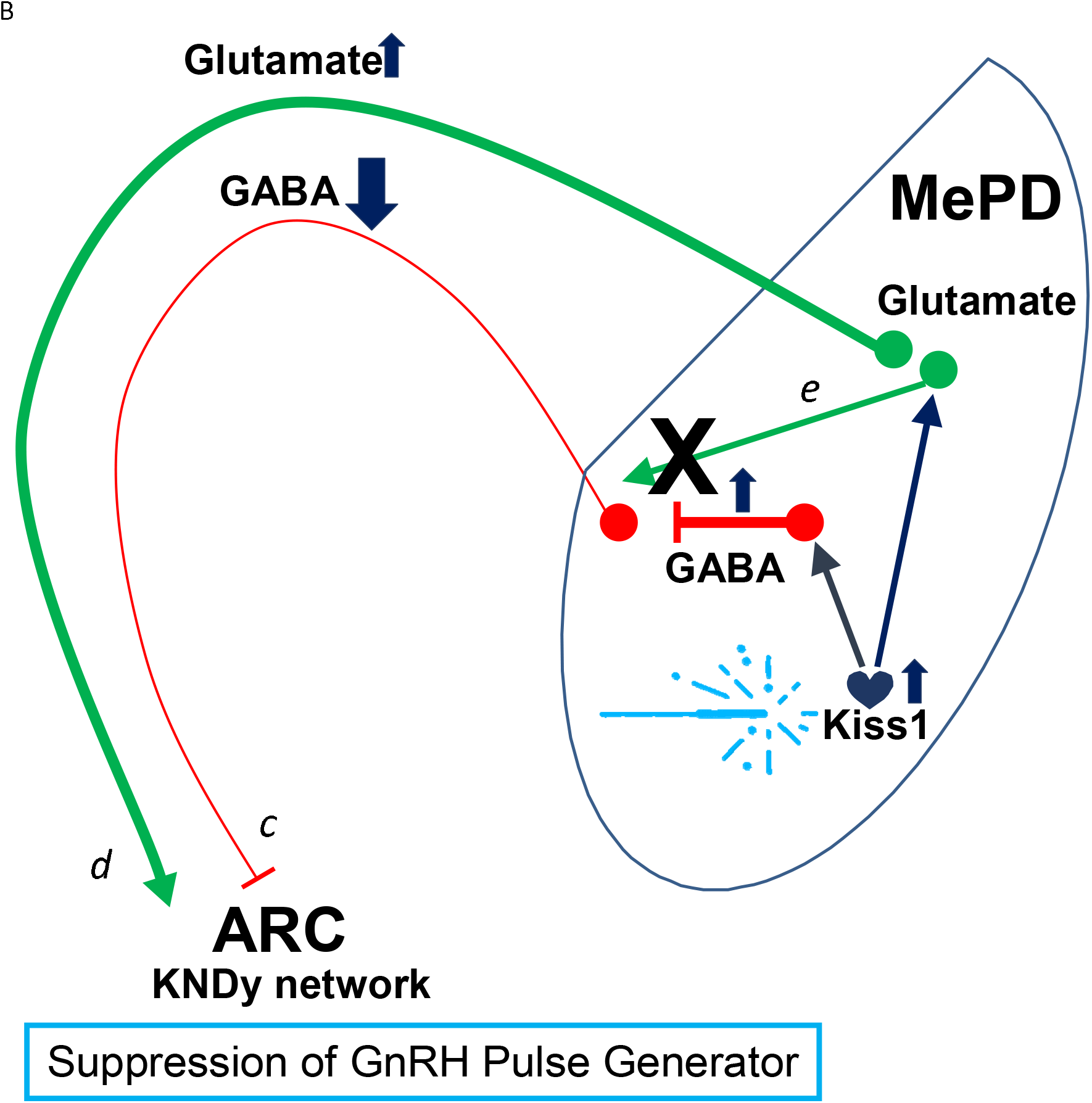
Model showing the proposed interactions and pathways involved in the MePD regulation over the hypothalamic GnRH pulse generator (ARC^KNDy^). (A) According to the hypothesis model, kisspeptin in the MePD regulates the GnRH pulse generator by activating two pathways: i) a GABA-GABA disinhibitory pathway and ii) a pathway involving glutamatergic MePD projection neurones. (B) Optogenetic stimulation of MePD Kiss1 (a) results in activation of the GABA-GABA disinhibitory pathway (b) that leads to a reduction in GABAergic tone arising from the MePD (c), as well as amplification of the MePD glutamatergic tone (d); it also causes activation of glutamatergic interneurones that project to the GABAergic efferent neurones, counteracting the stimulatory output of the MePD (e). Antagonism of these glutamatergic interneurones, combined with the optic stimulation of MePD Kiss1, results in the net effect of over-stimulation of the ARC KNDy network resulting in a transition from a pulsatile to a quiescent dynamic state of the GnRH pulse generator.

The fact that GABA_A_R antagonism alone failed to cause any change in LH pulse frequency is an important factor in our model: it is possible that under basal conditions MePD kisspeptin neurons are relatively quiescent, and therefore solely pharmacologically blocking the inputs to the GABAergic MePD projections without a corresponding increase in glutamatergic activity would make little change to the net influence of the MePD over the KNDy system. Although silent kisspeptin signalling under basal conditions supports the current hypothesised model, it contradicts neuropharmacology studies that show antagonising endogenous kisspeptin within the MePD causes a robust decrease in LH pulse frequency (Comninos *et al*., 2016). However, this latter study used OVX rats that were supplemented with 17 β-estradiol to mimic the hormonal profile in the diestrus phase of the estrous cycle. It is important to note, the present study used OVX mice which were not supplemented with 17 β-estradiol. *Kiss1* expression within the MePD varies in relation to the estrous cycle with low expression observed in OVX mice; estradiol treatment, however, amplified *Kiss1* expression in this brain region (Kim *et al*., 2011). This may explain why under basal conditions the MePD kisspeptin system appears reduced in our OVX mouse model.

Therefore, the proposed model does well to explain why GABA_A_R antagonism in the presence of MePD *Kiss1* optical stimulation reduces LH pulse frequency. However, a possible explanation as to why optogenetically stimulating these neurons in the presence of intra-MePD glutamate antagonists essentially stops all GnRH pulse generator activity is more complex. Complying with the model would suggest that blocking glutamate activity, while activating the GABA-GABA disinhibitory system, would in fact increase LH pulse frequency rather than prohibiting it altogether. Nevertheless, we provide a potential explanation for this phenomenon. By examining all of the individual pulse profiles more closely, a subtle, but potentially crucial aspect is identified. In over 80% (5 out of 6) of tests in which glutamate antagonists were infused in conjunction with optical stimulation, a pulse of LH was detected precisely 10 mins after the bolus infusion, and immediately *before* the onset of light stimulation. Only once the optic laser was switched on did the detection of LH pulses reliably cease. Therefore, it is reasonable to posit that while this protocol indeed blocked GnRH pulse generator activity, this occurs via a mechanism of potential over-stimulation which sends the KNDy system into a state of inertia as it is unable to respond. The proposed hypothesis is that activation of MePD kisspeptin drives i) the disinhibitory GABA-GABA pathway from the MePD, ii) glutamatergic interneurons that in turn project to the GABA-GABA pathway, and iii) glutamatergic projections from the MePD onto the ARC (a schematic diagram describing this is shown in Figure 3A). Thus, optogenetic stimulation of the MePD *Kiss1* neurons combined with antagonism of MePD glutamate results in the net effect of heightened activation of the GnRH pulse generator and resultant inertia (Figure 3B). The proposal of excessive GnRH pulse generator neuronal activity resulting in depolarisation silencing is in line with recent findings from our research group. Using mathematical models confirmed with *in vivo* optogenetics, it is now known that the ultradian oscillation of the hypothalamic KNDy network works on a bifurcation system that is eventually terminated as an upper threshold of basal neuronal activity is reached (Voliotis *et al*., 2019); we have described in detail how stimulation of the KNDy network increases in network excitability (e.g. via glutamatergic activity) or neurokinin B (NKB) signalling results in a robust transition from a pulsatile to a quiescent dynamic state of the GnRH pulse generator (Voliotis *et al*,. 2021). Furthermore, unpublished work conducted by O’Byrne using ovariectomised primates showed that administration of NMDA, a potent neuronal activator, evoked an MUA volley, the electrophysiological correlate of GnRH pulse generator activity (O’Byrne and Knobil, 1993) and corresponding LH pulse, followed by neuronal silence and cessation of LH pulses. Thus, the GnRH pulse generator is highly sensitive to incoming stimuli and may be prone to silencing by excessive activation. Importantly, glutamate antagonism alone did not result in a significant decrease in LH pulse frequency, and this is in line with the abovementioned theory of basal quiescence of the MePD kisspeptin system.

The present study also investigated the role of GABA_B_ signalling in the activity of MePD kisspeptin and the GnRH pulse generator using CGP-35348, a GABA_B_R selective antagonist. In contrast to the significant reduction in LH pulse frequency observed with bicuculline and optic stimulation, the interference of GABA_B_R signalling in conjunction with optogenetics only went so far as to prevent the increase in LH pulsatility, indicating a present, yet smaller, influence. The reason for this difference remains unclear, but can be potentially explained by the pharmacological differences between GABA_A_ and GABA_B_ receptors, with the former accounting for fast inhibition while the latter is responsible for slow inhibition (Nicoll *et al*., 1990). Moreover, it has been shown that in the case of LH release, only ICV activation of the GABA_A_R, and not GABA_B_R, both reduced LH release from the pituitary and GnRH levels in the POA (Leonhardt et al., 1995), suggesting a differential role for the two receptor subtypes in reproductive neuroendocrinology. Moreover, while it has been shown that knockout of the GABA_B_R subtype in adult female mice results in subfertile phenotypes such as decreased hypothalamic levels of GnRH and GnRH mRNA, it has no diminishing effects on LH or FSH levels, or *Kiss1* expression in the hypothalamus (Catalano *et al*., 2005; Catalano *et al*., 2010; di Giorgio *et al*., 2014); *Kiss1* levels in the MePD, however, are increased. This limited effect is in line with the present finding showing only partial consequences of MePD GABA_B_R antagonism together with kisspeptin optic stimulation.

These data have demonstrated, for the first time, the possible neuronal mechanisms by which increased kisspeptinergic activity within the amygdala increases GnRH pulse frequency, which could also provide the basis for the sexual development of puberty. It would be of interest to further investigate the roles of GABA and glutamate in this network, including in relation to perturbations of reproductive physiology associated with the amygdala such as stress and abnormal food intake.

## Acknowledgements

We are extremely grateful for all the help and advice on optogenetics provided by Dr. Matt Grubb, Centre for Developmental Neurobiology, Faculty of Life Sciences and Medicine, King’s College London, UK. The authors gratefully acknowledge the financial support from the MRC (MR/N022637/1) and BBSRC (BB/S000550/1). GL and DI are funded by MRC PhD studentships.

## Data Availability Statement

All data contained within the manuscript have been deposited in the King’s Research Data Management System and are freely available to public access (www.kcl.ac.uk/library/researchsupport/research-data-management/preserve/deposit-your-data-with-kings3).

## Notes

https://www.kcl.ac.uk/researchsupport

### Competing Interest Statement

The authors have declared no competing interest.

